# Osteoblast-Induced Collagen Alignment in a 3D *in vitro* Bone Model

**DOI:** 10.1101/2023.11.17.567552

**Authors:** Judith M. Schaart, Mariska Kea-te Lindert, Rona Roverts, Wouter H. Nijhuis, Nico Sommerdijk, Anat Akiva

**Affiliations:** Department of Medical BioSciences, Radboud University Medical Center, Geert Grooteplein Zuid 28, 6525GA Nijmegen, The Netherlands; Electron Microscopy Center, Radboudumc Technology Center Microscopy, Radboud University Medical Center, Geert Grooteplein Noord 29, 6525EZ Nijmegen, The Netherlands; Department of Orthopaedic Surgery, University Medical Centre Utrecht, Wilhelmina Children’s Hospital, 3508GA Utrecht, The Netherlands

**Keywords:** Bone formation, Collagen alignment, 3D cell culture, Volume electron microscopy, 3D FIB/SEM

## Abstract

The bone extracellular matrix consists of a highly organized collagen matrix that is mineralized by hydroxyapatite. Even though the structure and composition of bone have been studied extensively, the mechanisms underlying collagen matrix organization remain elusive. In this study, we developed a 3D cell culture system in which osteogenic cells deposit an oriented collagen matrix, that is subsequently mineralized. Using live fluorescence imaging combined with volume electron microscopy, we visualize the organization of the cells and collagen in the cell culture. We show that the osteogenic cells are organizing the collagen matrix during development. Based on the observation of tunnel-like structures surrounded by aligned collagen in the center of the culture, we propose that osteoblasts organize the deposited collagen during migration towards the periphery of the culture. Overall, we show that cell-matrix interactions are involved in collagen alignment during early-stage osteogenesis and that the matrix is organized by the osteoblasts in the absence of osteoclast activity.

## Introduction

Bone is a highly organized tissue of which the composition and organization of the extracellular matrix, that consists mainly of collagen type 1 mineralized with calcium phosphate, are the main determinants for its mechanical properties (Fratzl and Weinkamer, 2007; Weiner and Wagner, 1998). In mature bone, both in trabecular and in cortical osteonal bone, the collagen is arranged in bundles that, in turn, form ordered arrays embedded in a disordered material (Reznikov et al., 2014). However, in most species, including humans, the first bone to be deposited during skeletal development is woven bone, where the collagen has no preferred orientation (Weiner and Wagner, 1998). Other species, such as fast-growing juvenile mammals, are born with fibrolamellar bone, a form of transient primary bone tissue in which the collagen is highly aligned in one direction (Magal et al., 2014; Weiner and Wagner, 1998). During the lifetime of the organism, the extracellular matrix is remodeled through the interplay of the different bone cell types, in which osteoclasts resorb redundant bone material, osteoblasts produce new matrix where needed, and osteocytes orchestrate this interplay. This process is also responsible for the replacement of woven and fibrolamellar bone by the cortical and trabecular lamellar bone (Katsimbri, 2017; Magal et al., 2014; Weiner and Wagner, 1998).

Already in the late 80’s, Wagner and Weiner summarized the different 3D collagen organizations and their impact on the mechanical properties of the different bones, connecting structure to function (Weiner and Wagner, 1998). These models were later refined by Reznikov *et al*. (Buss et al., 2022; Reznikov et al., 2014) who used 3D focused ion beam scanning electron microscopy (3D FIB/SEM) to further elucidate the ultrastructural details of different bones. However, the mechanisms underlying the formation of these organized structures during bone formation remain largely unresolved.

In recent years, multiple 3D *in vitro* cell culture models for bone have been developed to study variable aspects of bone development (Akiva et al., 2021; Budyn et al., 2018; Nasello et al., 2020; Thrivikraman et al., 2019), which significantly improved the physiological relevance of bone cell cultures compared to previously used monolayer systems. These studies showed that mechanical stimulation (Akiva et al., 2021; Budyn et al., 2018) and matrix stiffness (Thrivikraman et al., 2019) are critical influencers of bone formation with regard to differentiation and bone matrix maturation. Remarkably, some of these 3D models showed that inducing longitudinal strain by fixation of a 3D cell culture on opposite sides was enough to induce matrix alignment and aligned mineralization (Iordachescu et al., 2018; Sasaki et al., 2010; Sasaki et al., 2015). As these models were based on only osteoblast and osteocyte-like cells, the parallel organization of the collagen cannot be attributed to a remodeling process involving osteoclasts. This suggests that such models could shed light on the mechanism of collagen orientation during fibrolamellar bone formation, which, although it occurs in many vertebrates, is still unknown.

In order to further analyze the processes underlying collagen matrix organization in the absence of osteoclastic remodeling, we developed a 3D osteogenic culture system containing a uniaxially fixed, degradable fibrin hydrogel, in which human mesenchymal stem cell (hMSC) derived osteoblasts could produce an aligned collagen matrix that was subsequently mineralized. We used this *in vitro* model to study the early-stage development of the aligned collagen matrix using fluorescence and 3D electron microscopy (volume EM). 3D FIB/SEM revealed the formation of long voids (tunnels) surrounded by layers of aligned collagen as a result of the migration of osteoblasts from the hypotrophic fibrin hydrogel core to the more nutrient-rich outer layers of the 3D culture. Formation of these “tunnels” requires a high level of control over the collagen deposition (and removal) by the osteoblasts, and we propose that a similar mechanism is likely active during fibrolamellar bone formation in many vertebrates.

## Results

### Establishing & Characterizing a 3D Bone Culture

A uniaxially fixed 3D cell culture was used to study cellular and collagen organization in developing bone. hMSCs were seeded as a homogeneous suspension in fibrin, supported by Velcro® strips on each side to introduce strain (Figure 1A-B). This setting facilitates cell stretching and alignment of various cell types in a 3D environment (de Jonge et al., 2013; Iordachescu et al., 2018). Two days after seeding, the culture medium was supplemented with dexamethasone and β-glycerophosphate to induce osteogenic differentiation (Jaiswal et al., 1998; Tenenbaum, 1981; Tenenbaum and Heersche, 1982). To analyze the osteogenic differentiation and architecture of the culture, 3D fluorescence microscopy and scanning electron microscopy (SEM) were combined with biochemical assays (Figure 1C-F).

**Figure 1.**
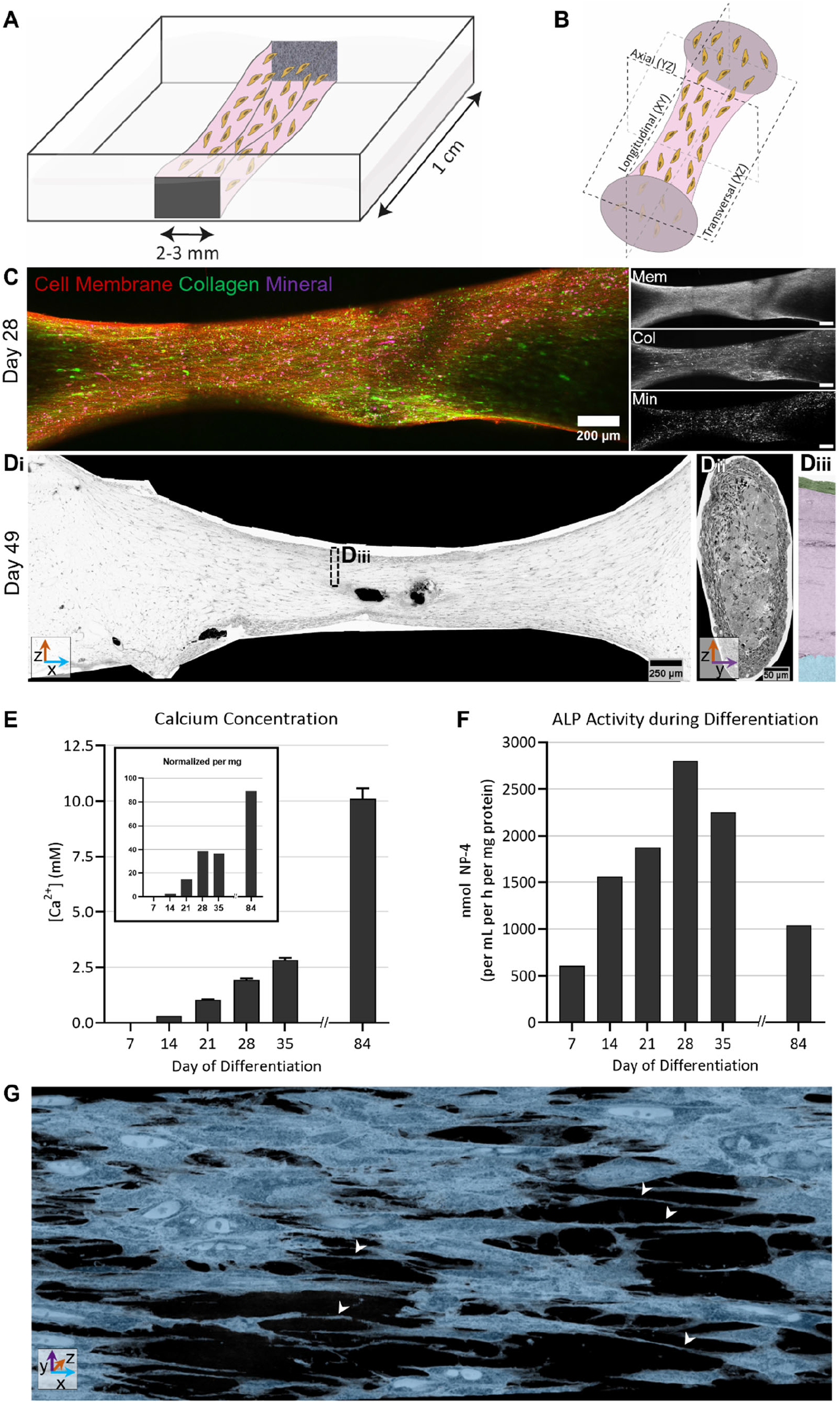
Establishing a 3D osteogenic cell culture to study collagen organization. A) Schematic representation of the fibrin/cell suspension (pink) between two Velcro® fixation points (grey, 2-3mm width). The culture is grown in an Ibidi μ-slide 8-well slide, and has a length of 1 cm. B) Schematic representation of the image planes within the 3D cell culture. Transversal (XZ), longitudinal (XY) and axial (YZ) views were obtained using 3D imaging approaches. C) The cellular distribution (Cell Mask Deep Red, red), collagen production (CNA35-OG488, green), and mineral deposition (Calcein blue, magenta) as visualized by maximum intensity projections of overview fluorescence images at 28 days of differentiation. Single channels in gray. Scale bars: 200 μm. D) SEM overview of the cross-sections of a heavy metal stained and embedded culture at day 49 of differentiation showing the outline and general organization of the osteogenic culture along the longitudinal axis (i, scale bar: 250 μm) and axial axis (ii, scale bar: 50 μm). 3 regions can be identified (iii) based on the presence of high cell density (green), mainly collagen (magenta) and fibrin (blue). E) Quantification of the deposited calcium in the matrix showed an increasing mineral content during the differentiation period. After normalization of the calcium content to the dry weight (mg) of the culture, a similar trend was observed (inset). F) Quantification of the ALP activity (normalized to the protein quantity in the sample as determined by Bradford assay, Supplementary Figure A.1D), to determine osteogenic differentiation status. G) 3D reconstruction of segmented cells from 3D AT-SEM in the fibrin-rich core of the culture. Cells show multiple protrusions 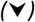 and intercellular connections.

The cellular distribution, collagen production, and mineral deposition were examined over the course of 28 days using live fluorescence microscopy (Figure 1C, Supplementary Figure A.1A). Evenly distributed cells and collagen, both aligned along the longitudinal direction, were present after 7 days of differentiation, indicating that, at this time, osteogenesis had already started. The first mineral signal was visible after 21 days of differentiation (Supplementary Figure A.1A), and the mineral was clearly present at day 28 (Figure 1C). During differentiation, the degradation of the fibrin scaffold was actively controlled by removing amino-caproic acid (de Jonge et al., 2013), an inhibitor of the fibrinolytic plasminogen, from the culture medium after 14 days of differentiation, after a sufficiently dense collagen matrix had been deposited by the cells. In addition, live/dead staining demonstrated the viability of the majority of the cell culture after 21 days of differentiation (Supplementary Figure A.1B).

Since the probing depth of the fluorescence imaging was limited by the high density of the cell culture (Supplementary Figure A.1C), the transversal and axial planes of the culture were imaged by SEM to visualize the composition of the bulk of the culture (Figure 1D_i_-D_ii_). These overviews showed a structure similar to the maximum intensity projections of the fluorescence imaging (Figure 1C), where the edges of the culture that interface with the Velcro® are significantly wider than the central region. This is due to the limited contraction of the gel in the region that interacts with the Velcro® anchors, which have a width of 2 to 3 mm. In combination with the thin axial cross-section of the culture (Figure 1D_ii_), we concluded on a symmetrical radial structure of the culture. Based on the longitudinal overview, we further identified 3 distinct regions for further analysis, from the peripherical area towards the core (Figure 1D_iii_): (i) cell-dense layer (periphery, indicated in green), (ii) collagen-rich area (bulk, indicated in magenta), and (iii) fibrin-rich area (core, indicated in blue).

In addition to these image-based analyses, biochemical assays were performed to confirm osteogenesis by quantifying calcium concentration and alkaline phosphatase (ALP) activity (Figure 1E,F). Quantification of calcium deposition in the matrix showed an overall increasing mineral content during the differentiation (Figure 1E), and this trend was retained after normalization of the calcium content to the dry weight of the demineralized cultures (Figure 1E, inset).

The cellular differentiation status in the mineralizing culture was then analyzed by quantification of ALP activity normalized to the total protein content (Figure 1F, Supplementary Figure A.1D). As expected (Husch et al., 2023; Meesuk et al., 2022; Westhauser et al., 2019), the ALP activity increased during differentiation from day 7 to day 28, after which it decreased again. The increasing ALP activity indicated that the hMSCs were maturing towards osteoblasts, as ALP expression is a marker for early osteoblast activity (Golub and Boesze-Battaglia, 2007; Zernik et al., 1990). The subsequent decrease could indicate the differentiation of osteoblasts towards more mature post-osteoblast or pre-osteocyte stages (Miron and Zhang, 2012).

Subsequently, the morphology of the cells in the fibrin-rich core of the culture was investigated by 3D array tomography SEM (AT-SEM), to analyze the cell content. This showed that most of the cells in the core of the culture have a healthy appearance despite their distance (∼250-300 μm) from the nutrient-rich medium at the surface (Supplementary Figure A.1E). We inferred their healthy appearance from their size and shape and the morphology of their cellular organelles (e.g. mitochondria, nuclei, and endoplasmic reticulum). A deep-learning-based semantic segmentation algorithm was trained to automatically annotate the cells in this 3D image stack, which enabled the visualization of the 3-dimensional environment and cellular interactions. The segmentation revealed an interconnected network of cells, showing multiple protrusions and interactions (Figure 1G, Supplementary Video 1). This morphology is characteristic for osteocyte-like cells, and in line with the hypothesis that part of the osteoblasts could already have differentiated towards the final stage of osteogenesis. Overall, these data confirmed that the culture resembled crucial characteristics of bone development, including collagen production and mineralization.

### Cellular and Collagen Organization during Osteogenic Differentiation

To further analyze the cell and collagen organization during osteogenic differentiation in a dynamic approach and from an early stage of collagen production, live fluorescence microscopy was used to weekly analyze the orientation of these components in living cell cultures (Figure 2). To visualize the alignment of cells and collagen in the periphery of the culture, a color-based orientation analysis was performed on Z-stack maximum intensity projections of differentiating cultures (Figure 2A). This showed that most of the visible cells aligned along the direction of attachment (∼90°) from day 7 onwards and that the cellular orientation remained stable over time. Moreover, the deposited collagen was oriented mainly in the same direction as the cells. Corresponding directionality histograms confirmed the alignment of the cells and collagen in a direction parallel to the attachment in all stages of differentiation (Figure 2B). However, overlaying the orientation histograms of the cells (Figure 2B, red) and collagen (Figure 2B, green) in the cultures at multiple differentiation time points showed that the orientation of these two components was not perfectly overlapping. When the main orientation of the collagen and the cells were compared, it was observed that the angle between these two components increased from day 4 to day 35 of differentiation (Figure 2C). After this time point, the angle between the major orientations showed a decreasing trend.

**Figure 2.**
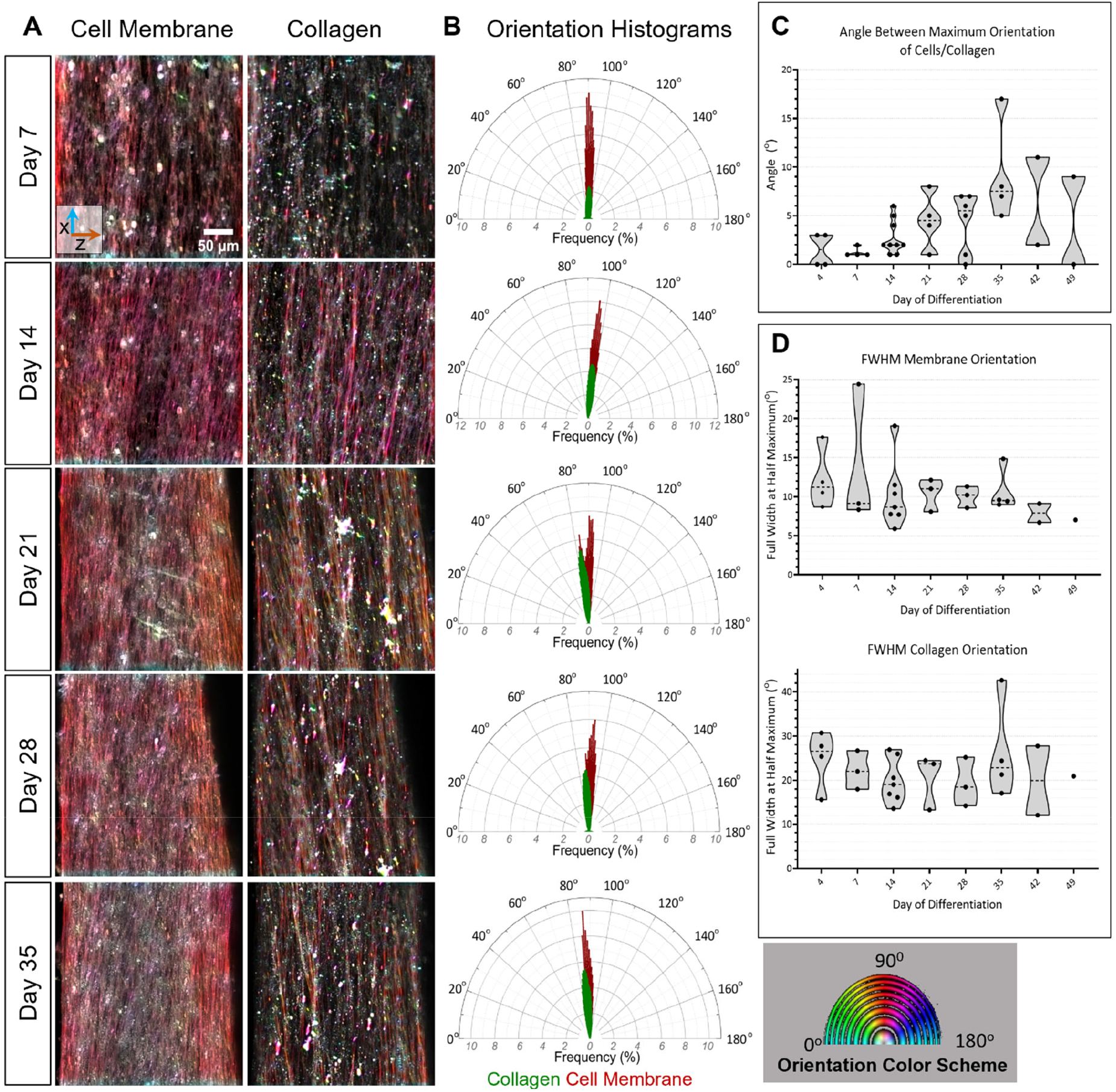
Orientation of cells and collagen during differentiation. A) Color-based orientation analysis of fluorescence images (maximum intensity projections) of the cell membranes (left) and collagen matrix (right) showing that most of the visible cells aligned along the direction of the fixation. Scalebar: 50 μm. B) Orientation histograms corresponding to the fluorescence images show that the collagen deposited by the cells was oriented mainly in one direction. The plots indicated a small offset between the orientation of the cells and collagen. C) The main orientation of the collagen and the cells in multiple experiments were compared, and the angle between the two maxima was determined and plotted over time. The data indicate that the angle between these two components increased over time from day 7 to day 35 of differentiation. The angle decreased again after this time point. D) The FWHM of the directionality histograms was calculated by Lorentzian fitting and plotted for different differentiation times. The FWHM of the cell alignment was overall smaller than the FWHM of the collagen alignment during the complete culture period. The FWHM of both cellular and collagen orientation showed a slightly decreasing trend during the time of differentiation.

Subsequently, the alignment quality over time was analyzed by calculating the full width at half maximum (FWHM) of the directionality histograms by Lorentzian fitting (Figure 2D). Overall, the FWHM of the cell alignment (7-14°) was smaller than the FWHM of the collagen alignment (19-26°), indicating that the cells aligned more uniformly than the collagen. Furthermore, the FWHM of both components was quite stable, indicating that the alignment quality of the cells and the collagen did not change significantly during culture development. The fact that the cellular organization did not change over time indicates that the cellular morphology is stable at an osteoblast-like stage and in this time-frame does not convert during differentiation to an osteocyte-like phenotype that is expected to have more protrusions. However, the morphology of the cells observed in these fluorescence microscopy experiments only represents the cells in the periphery of the culture, as the bulk and core regions of the culture cannot be visualized due to the limited penetration depth of the live fluorescence imaging.

### Cell-Collagen Interfaces in the Bulk and Core Regions

To further visualize and analyze the development of the collagen and the cell-collagen interactions, 3D volume EM (FIB/SEM) was performed on three regions of interest. These regions were located on the border of the cell-rich periphery and collagen-rich bulk, as well as on the border between the bulk and the fibrin-rich region. Using these stacks, we then further analyzed the development of the different collagen-rich regions to evaluate collagen alignment, packing, and orientation in space (Figure 3, Supplementary Video 2-5). Variable densities of collagen were observed, dependent on the location in the culture e.g. close to the periphery (higher density) or close to the core (lower density, Supplementary Table A.1).

**Figure 3.**
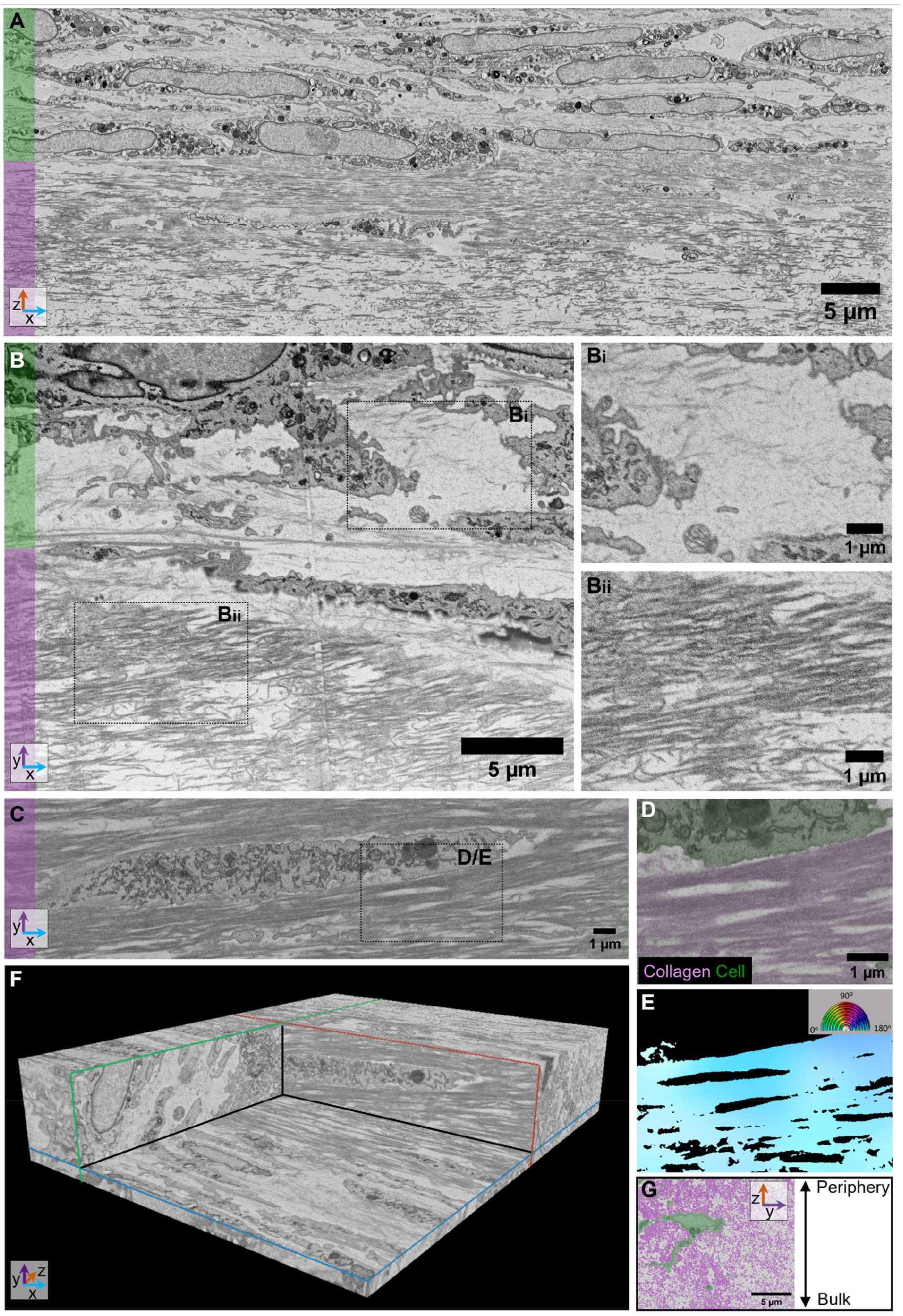
Cell-collagen interface in the periphery and bulk region. A) Resliced transversal view from 3D FIB/SEM stack on the border of the periphery and the bulk region. The periphery region contains a high density of cells, which borders a dense region of highly aligned collagen with lower cell density in the bulk region. Scalebar: 5 μm. B) Longitudinal view from FIB/SEM imaging at the border of the periphery and the bulk region shows different collagen organizations in the different regions (scalebar: 5 μm). Collagen in the cell-dense region is observed as thin, disorganized fibers (B_i_, scalebar: 1 μm), while the first layers of the bulk region consist of thicker fibers with higher density and organization (B_ii_, scalebar: 1 μm). C) FIB/SEM image from a cell embedded in collagen in the bulk region. Scalebar: 1 μm. D) Segmented cell (green) and collagen from ROI from C. E) Color-based orientation map from collagen segmentation (ROI from C). Scalebar: 1 μm. F) Cross-sections from 3D reconstruction of FIB/SEM region, showing a cell surrounded by dense and aligned collagen. On the axial plane (YZ, green) collagen is visible as dots, while on the longitudinal (XY, red) and transversal (XZ, blue) planes the collagen is seen as threads. G) Axial view of a cell (segmented in green) with a body and tail, proposedly migrating through the collagen from the bulk towards the periphery region. Scalebar: 5 μm.

Resliced views of the transversal plane on the border of the periphery and bulk region (Figure 3A) showed that the peripheral region consisted of a layer of tightly packed cells with a thickness of three to five cellular layers. Surprisingly, most cells showed an elongated structure rather than a cubic one that is more phenotypic of active osteoblasts (Blair et al., 2017). However, the high amount of organelles, including the mitochondrial network, that the cells show suggests that these are still very active cells (Blair et al., 2017; Kassem et al., 1992). The cellular layer lines a relatively thick collagen layer of about 5 μm, in which the collagen is dense and highly aligned (Figure 3A-B, magenta). In this region, we can define a clear borderline between the cell-dense region and the aligned collagen (Figure 3B), along which the cells stretched with rather long protrusions in the direction of the strain. In between the stretched cells, the low amount of collagen showed random orientations (Figure 3B_i_), and the collagen fibers seemed thinner and shorter than in the highly aligned collagen layer bordering the cell layer (Figure 3B_ii_). This randomly organized collagen is, most likely, newly produced collagen that has not been aligned yet.

To visualize the relation between the cells and the collagen in the 3D volume of the highly dense collagen, deep-learning-based segmentation of collagen fibrils and cells in 3D was performed on a selected region of the image stack (Figure 3C-D). The 3D visualization (Figure 3F, Supplementary Video 2) showed that cells in the bulk region were completely embedded in the dense, well-aligned collagen, and even though in this area the observed cell density was low, the cells still aligned in the direction of the strain. Subsequently, binary images of the segmented collagen were used for orientation analysis (Figure 3D-E), which confirmed the high level of alignment in this region. This alignment was present in all frames throughout the complete stack (Supplementary Figure A.2A). Even in the core region of the culture, where the density of the collagen was relatively low, some degree of orientation was still observed (Supplementary Figure A.2B, Supplementary Video 3-4). Combining the information from the longitudinal and axial image planes revealed that near the periphery the collagen directly surrounding the embedded cells closely followed the outline of the cell (Figure 3C,F).

Interestingly, the resliced axial views of the periphery/bulk stack showed the presence of cells with elongated cell protrusions towards the core of the culture and not parallel to the strain direction (Figure 3G, Supplementary Figure A.3), which had morphologies typical for migrating cells, in which cellular protrusions are elongating from the cell body in the direction of migration (Lu and Lu, 2021). These cells were frequently in close proximity to regions in which no cellular structures or matrix were observed. From this we hypothesized that these cells were migrating towards the nutrient-rich peripheral region, using these spaces to relocate their cell body, while interacting with the collagen to organize and condense the matrix.

### Cell-Shaped Tunnels and Voids in the Collagen Matrix

Next, the collagen organization and cell-collagen interfaces in the FIB/SEM stack recorded around the bulk/fibrin border were analyzed (Figure 4). Overall, the collagen density in this region was lower than close to the periphery (Supplementary Table A.1). However, more condensed regions of well-aligned collagen were observed in close proximity to cells (Figure 4A-C), and cells still show a preferred orientation along the longitudinal axis of the culture. In addition to the cells embedded in denser and well-aligned collagen, cell-free regions surrounded by organized collagen extending from cell protrusions were also observed (Figure 4A, *). Interestingly, although cells are not detected in these regions, these collagen fibrils border voids that could fit cell protrusions. After automated segmentation of cells and collagen, the 3D reconstruction of the cell protrusions and the proximate collagen showed that these cell-shaped voids form tunnels, outlined by collagen walls (Figure 4B, collagen: magenta, cell: green, Supplementary Video 5), which is also confirmed by visualization of the axial view of these voids (Figure 4B_i_).

**Figure 4.**
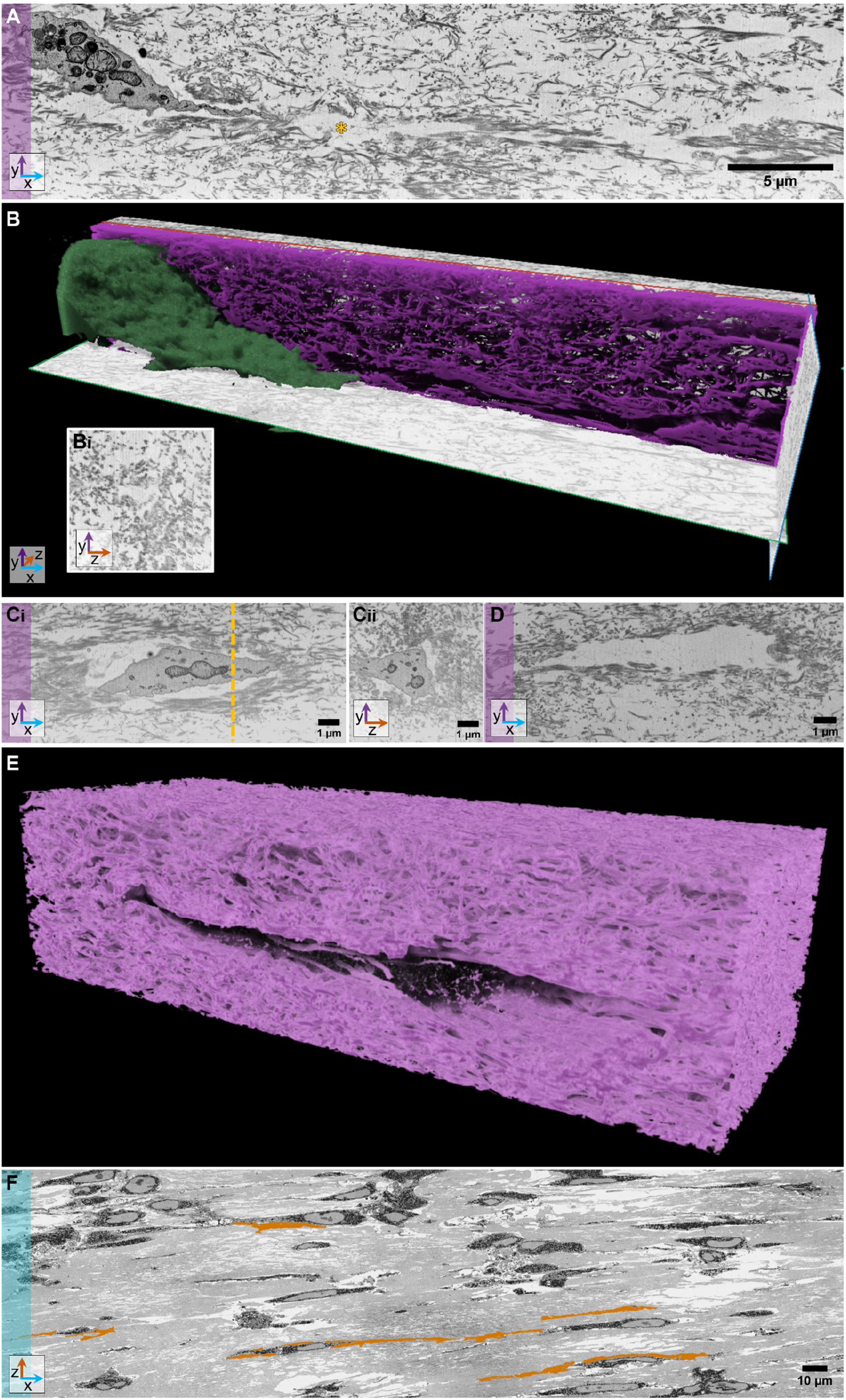
Tunnels and voids surrounded by collagen indicate active collagen organization by osteoblasts. A) FIB/SEM images along the longitudinal (XY) axis showed the presence of regions in the cell culture where the collagen aligned around a cell-free space extending from a cell protrusion (*). B) 3D reconstruction of the segmented cell (green) and collagen (magenta) in the 3D FIB/SEM stack showed that the cell-free space was shaped as a tunnel. Reslicing of the axial view (B_i_) around the tunnel, showed that the tunnel was surrounded by collagen which showed a decreasing density at a larger distance from the center of the tunnel. C) FIB/SEM imaging (longitudinal, XY) showed that collagen in these cultures had a high density and alignment in close proximity to cells (C_i_). The resliced axial view (C_ii_, YZ, at dotted line) showed a gradient in collagen density around the cell. Scalebars: 1 μm. D) FIB/SEM imaging showed cell-free voids surrounded by dense and aligned collagen that showed a gradient when moving further away from the cell. Scalebar: 1 μm. E) 3D reconstruction of segmented collagen (magenta) surrounding a cell-shaped void region. F) AT-SEM imaging in the fibrin-rich core region showed the presence of voids (orange) next to cells in the mixed fibrin/collagen matrix. Scalebar: 10 μm.

When comparing the collagen density around these tunnels (Figure 4A-B_i_) to the collagen density around a cell body (Figure 4C) it was observed that highly aligned collagen is present both around the cells (Figure 4C) and the voids (Figure 4A). Additionally, the axial cross-sections show a gradient in the collagen density surrounding the cells and the voids, with a pattern of more densely packed and better-organized collagen closer to the cells (Figure 4B_i_) and voids (Figure 4C_ii_). We therefore propose that these cell-free tunnels are created by the retraction of the cell after depositing the aligned collagen layer.

Similar observations regarding collagen organization and density were made for regions with collagen surrounded cell-shaped voids (Figure 4D). Even though no neighboring cell was observed in the reconstructed volume, the cell-like shape remained as a hole in the aligned collagen matrix (Figure 4E).

When analyzing the fibrin-rich core of the culture, 3D AT-SEM showed that cells still have a preferred alignment along the longitudinal axis of the culture (Figure 4F). Interestingly, also the fibrin-embedded cells were regularly surrounded by regions in which no cellular structures or protein matrix were observed in the 3D AT-SEM volume, and the tunnels seemed to form connections between different cells (Figure 4F, orange, Supplementary Figure A.2C). Looking at these data in more detail showed that the observed tunnels were more tight in regions with a mixed fibrin-collagen matrix (Supplementary Figure A.2D, *), while the voids of cells surrounded by pure fibrin were wider and less structured (Supplementary Figure A.2D, ▾). This could possibly be due to higher stability induced by crosslinking of the *collagen* matrix, as well as degradation of the *fibrin* matrix by the cells. Overall, these observations indicated that the extracellular matrix in these cultures was organized during osteogenic differentiation by osteogenic cells in the culture.

## Discussion

In this study we developed a human MSC-based 3D cell culture using uniaxially fixed fibrin matrices, in which osteogenic differentiation, collagen matrix development, and the collagen-cell interface were studied. In accordance with previous reports (Iordachescu et al., 2018; Kim et al., 2021; Matsumoto et al., 2007; Sasaki et al., 2010; Sasaki et al., 2015), we observed that the uniaxial fixation of the culture induced alignment of the cells and the deposited collagen matrix solely through the activity of osteoblasts and osteocyte-like cells. Using a 3D multiscale imaging approach, combining 3D array tomography SEM, live fluorescence imaging, and 3D FIB/SEM (Figure 5), we were able to reveal structural details of the resulting culture that allow us to propose that the alignment of the newly deposited collagen matrix derives from the migration of the osteogenic cells as detailed below.

**Figure 5.**
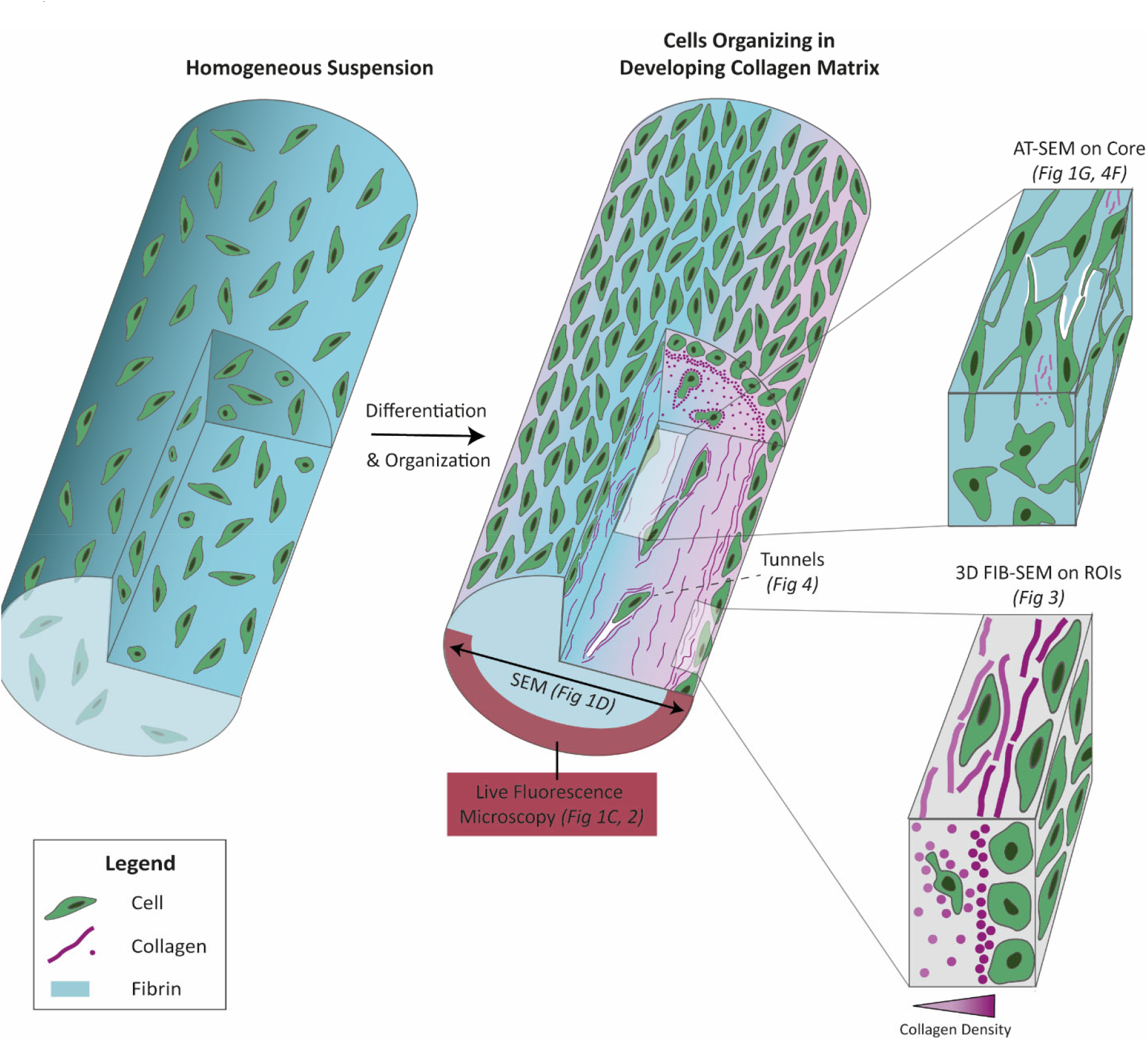
Schematic representation of the development and multiscale analysis of our 3D osteogenic culture. At the moment of seeding, cells are randomly distributed through the fibrin in a homogenous suspension. After gelation and differentiation, the cells have reorganized into a radial structure with a dense layer of aligned cells on the periphery surrounding a dense layer of collagen, which can be observed by live fluorescence microscopy. SEM, AT-SEM, and 3D FIB/SEM allowed for the reconstruction of the 3D environment of the periphery as well as the bulk of the culture, showing that the collagen and cell density decrease towards the bulk of the culture, while the cells present in this region are surrounded by denser collagen. This indicates that the cells migrate throughout the culture towards the periphery, during which the collagen is contracted and organized.

In addition to highly aligned layers of collagen that surround the embedded cells and follow their surface, collagen *tunnels* were observed in the bulk of the culture. In the latter, the collagen appears to have followed the shape of a cell or cellular protrusion, although these cellular structures were no longer observed. Here, the cell seems to have retracted after alignment or even to have completely moved out (Figure 4). The fact that well-defined tunnels are observed in the regions of high collagen density, while those in the fibrin-rich regions were much less well defined, suggests that the cross-linking of the collagen plays a role in the persistence of their shape after cells migrate away (Rodriguez-Pascual and Slatter, 2016). Similar stable collagen structures have been observed in studies using fibroblasts (Guidry and Grinnell, 1987) and fibrosarcoma cells (Fisher et al., 2009). However, their role and dynamics have not yet been elucidated.

Our data show that in our osteogenic culture, the density of the collagen was not a determining factor for fibril alignment, as both the high-density collagen in the peripheral region of the culture and the collagen in the lower-density bulk and core regions showed alignment. However, the collagen showed higher degrees of alignment in regions where cells are present than in the surrounding areas, suggesting that the osteoblasts play an active role in collagen organization.

Moreover, the observation of collagen with lower degrees of alignment closer to the fibrin-rich core of the culture as well as in between the dense cells in the peripheral layer, together with highly aligned collagen around cells (Figure 3, Figure 4 & Supplementary Figure A.2) leads us to hypothesize that during osteogenesis collagen is quickly deposited in a disorganized fashion, resembling the structures observed in woven bone (Weiner and Wagner, 1998), and that later longer cell-collagen interactions induce collagen alignment and organization. This phenomenon of cells actively re-arranging collagen after deposition has been previously observed in monolayer cultures of mouse osteoblasts (Lu et al., 2018) and in 3D cultures for other cell types, such as cancer cells (Kim et al., 2018; Miller et al., 2020; Pamonag et al., 2022). The re-arrangement of collagen by cancer cells is essential to mediate migration and thus invasion (Kim et al., 2018; Pamonag et al., 2022), while for the osteoblasts in our cultures, it is essential to develop the cell-cell interactions that are critical for osteogenic development (Kim and Adachi, 2019; Stains and Civitelli, 2005).

We observe a clearly distinguishable border between a cell-dense layer and a collagen-dense layer, even though at the start of the experiment, the cells were randomly and homogeneously distributed in the fibrin gel, indicating that the cells move through the deposited collagen during differentiation. The 3D reconstructions of the cells embedded in the dense collagen (Supplementary Figure A.3) suggest that the cells move towards the nutrient-rich periphery of the culture, while also the alignment of the collagen in lower-density areas points to the movement of the cells towards nutritional and oxygen gradients. The cell-matrix interactions during these movements will be associated with shear stress and traction forces (Lange and Fabry, 2013) and are likely the cause of the collagen organization. We propose that the high degree of alignment in the high-density collagen layers is caused by the repeated interaction with moving cells and the compacting of the packed fibrils by the dense, aligned cell layer.

Here, the interaction with osteoblasts leading to collagen alignment could be mediated by collagen binding integrins, such as integrin α_2_β_1_ (Boraschi-Diaz et al., 2017; Kundu et al., 2009; Lee et al., 2020), of which the expression is increased during early osteogenic differentiation of hMSCs, followed by a decrease in later stages (Kundu et al., 2009). On the other hand, osteoblasts could potentially modulate their collagen-binding capacity by expressing non-integrin-dependent collagen-binding proteins, such as discoidin domain receptors (DDR) (Yang et al., 2020), which were shown to influence collagen patterning during bone development in early stage embryogenesis (Vogel et al., 1997).

## Conclusion

The 3D multiscale analysis of the developing uniaxially fixed human osteogenic cultures showed us variable levels of cell and collagen alignment in different regions of the culture. We propose that in the early phase of osteogenesis, the cells quickly deposit collagen with a low degree of organization Based on the fact that we start the culture from a homogenous suspension of cells in fibrin and the final organization that we observed, we further propose that during differentiation, osteoblasts move through the matrix towards the peripheral region of the culture, organizing the collagen they come in contact with and leaving tunnels in the matrix after they have moved on. We observe that tunnels in collagen-rich regions appear more shape-persistent than tunnels in fibrin-rich regions, most probably due to stabilization by collagen cross-linking. Cell-dense layers show a high degree of collagen alignment, supporting the active role of the cells that must impose shear forces on the matrix during their movement. With this, we show that cell-matrix interactions play a crucial role in collagen alignment in early-stage osteogenesis, and that, in particular, cellular motion can generate organized lamellar collagen matrices that subsequently can be mineralized.

In most vertebrates, including humans, the first matrix deposited during osteogenesis is disorganized woven bone that is only after remodeling by the interplay of osteoblasts and osteoclasts converted into osteonal bone with organized layers of mineralized lamellar collagen (Katsimbri, 2017; Weiner and Wagner, 1998). Indeed, we recently showed that a human organoid grown under non-directional mechanical stimulation produced disorganized, woven bone (Akiva et al., 2021). In contrast, fibrolamellar bone, the first deposited form of bone in many other, non-human, vertebrates, already shows a high degree of parallel collagen alignment prior to remodeling (Magal et al., 2014; Weiner and Wagner, 1998). Our experiments, and those of others (Iordachescu et al., 2018; Sasaki et al., 2010; Sasaki et al., 2015), show that the alignment of collagen can also occur in the early stages of human osteogenesis and hence is not genetically directed nor specific to species producing fibrolamellar bone. Instead, it can be induced by applying longitudinal stress.

Our results thereby contribute to understanding the mechanistic aspects underlying collagen organization in fibrolamellar bone, a topic that is under-investigated despite its importance for many species. Moreover, the ability to organize collagen in 3D osteogenic cell cultures without osteoclastic remodeling may provide new routes for the tissue engineering of load-bearing bone, as it has been notoriously difficult to achieve mechanically robust, organized bone matrices through an interplay of osteoblasts and osteoclasts in *in vitro* cell cultures (de Wildt et al., 2019).

## Materials and Methods

Chemicals and cell culture supplements were bought from Sigma-Aldrich/Merck unless indicated differently. Cell culture media and antibiotics were bought from Gibco.

### Cell Culture

Human mesenchymal stem cells (hMSCs) were isolated from bone marrow in University Medical Center Utrecht as described previously (Pennings et al., 2019). In short, human bone plus marrow waste material from a 11-year old female was obtained in accordance with the Declaration of Helsinki, with the approval of the Assessment Committee Biobank (TCBio) of the University Medical Center Utrecht (Utrecht, The Netherlands) under the approved number TCBio-18-685, and with the written consent of the parents. Bone marrow was diluted with an equal volume of phosphate buffered saline (PBS) and filtered through a cell strainer (100μm). Mononuclear cells (MNC) were isolated by centrifugation (320g, 20 min, RT) on a Ficoll-Paque gradient. The MNCs were washed with PBS to remove the Ficoll and seeded at a density of 250.000 cells per cm^2^ in MSC starting medium (αMEM (12571), 10% FCS, 100 U/mL penicillin/100 μg/mL streptomycin (PenStrep, 15140122), 0.2mM ascorbic acid and 1 ng/mL recombinant human FGF (R&D system, 233-GMP-025). The adherent cells were split by trypsinization at 80% confluency and frozen at passage 2 (50% αMEM, 40% FCS, 10% DMSO) in a Mr. Frosty (overnight). After freezing, the vials were stored in LN_2_.

hMSCs (passage 2 until passage 9) were cultured in a humidified incubator (37°C, 5% CO_2_) in MSC proliferation medium (PM; αMEM, 10% FCS, 100 U/mL-100 μg/mL PenStrep, 0.28 mM l-ascorbic acid-2-phosphate (AA2P (49752)), 1 ng/mL recombinant human FGF. Upon 80% confluency, cells were split (1:10) by trypsinization (0.05% Trypsin/0.53mM EDTA in PBS) according to standard cell culture protocols.

### Aligned 3D culture supported by Velcro®

The 3D cell cultures were grown in manually customized Ibidi μ-slide 8-wells slides (uncoated, 81501). The top edge of the μ-slides was lowered with an electrical polisher, after which the wells were cleaned in ethanol (70%) in an ultrasonic bath (5 min). Before use, the equipment and components were sterilized in 70% ethanol. Pieces of Velcro® (∼2x2mm, soft side) were attached to the edge of the wells with vacuum grease, while making sure that the hairs were facing each other (Figure 1A). Then, polystyrene walls covered with a thin layer of vacuum grease on the edges were placed on both sides of the Velcro® to create a closed cavity in the middle of the well in which the cell suspension was sequentially seeded.

Stock solutions of bovine thrombin (605157, 50U/mL in αMEM, sterilized on a 0.2μm filter) and bovine fibrinogen (60 mg/mL in αMEM, based on 77% clottable protein (F8630) were prepared in a laminar flow cabinet. Aliquots were thawed on ice and thrombin was diluted in PM to a concentration of 5U/mL. Fibrinogen was diluted to a concentration of 6 mg/mL in PM supplemented with amino-caproic acid (ACA, 2 mg/mL (A7824)).

Cells were trypsinized, counted, pelleted and resuspended in the diluted thrombin solution (2x10^6^ cells/mL thrombin). Next, the fibrinogen solution was mixed with an equal volume of cell suspension (final concentrations: thrombin 2.5U/mL, fibrinogen 3 mg/mL, ACA 1 mg/mL, cells: 1x10^6^/mL) and 100 μL of the mix was pipetted between the walls in each well, making sure that the suspension touched the Velcro®. The cells were polymerized (1h, 37°C, 5% CO_2_) before the polystyrene walls were removed with help of a scalpel. Next, the cultures were grown for 2 days in PM supplemented with ACA (1 mg/mL, 300 μL per well).

After 2 days, osteogenic differentiation was started by replacing PM with osteogenic induction medium (OIM; PM supplemented with 10 mM β-Glycerophosphate (βGP, G9422) and 100 nM dexamethasone (D4902)). Medium was refreshed three times a week. During the first 14 days of differentiation, the OIM was supplemented with ACA (1 mg/mL), while afterward, ACA was removed.

### Calcium assay

Cultures were harvested between 7 and 84 days of differentiation. The cultures were removed from the wells and washed in PBS. Then, the culture was sliced into three pieces using a scalpel, and one part was incubated in HCl (0.6M, 200μL) for 6 days (RT) to extract the mineral from the cultures, after which the samples were stored at -20°C until use. For quantitative analysis of the calcium content, the samples were thawed on ice, and the supernatant was used for an o-cresolphtalein colorimetric assay (Randox), according to the manufacturer’s instructions. In short, the reagents A and B were mixed in equal volumes and 250 μL was added to the samples (20 μL, pure or 10x diluted in MQ). The reaction was incubated (15 min, RT) and the absorption was measured using a platereader (570 nm, Biorad Benchmark Plus). Afterwards, the solid matrix was removed from the HCl and air-dried. The solids were weighted to use for normalization of the calcium concentrations.

### ALP activity assay

To determine the ALP activity of the cell cultures, one third of each gel was stored in MQ (200 μL, -20°C). The sample was lysed by 3 sequential freeze-thaw cycles and the solids were pelleted (1500g, 5 min). The supernatant was used for an ALP activity assay, based on para-nitrophenyl phosphate (pNPP, S0942) conversion to 4-nitrophenol (241326). The cell lysates (50 μL diluted in 50 μL Alkaline Buffer Solution (0.5M, Sigma, A9226)) were incubated with pNPP (100 μL, 5 mM in Alkaline Buffer) for 60 min at 37°C before the reaction was stopped by adding NaOH (0.3M, 100 μL). Then, the fluorescence was determined using a Victor Fluorescence Platereader (405nm, Biorad).

To normalize the activity, the samples were analyzed in a Bradford microplate assay (Biorad, 500-0006), according to the manufacturer’s protocol. In short, 10 μL of sample was mixed with 200 uL bradford reagent (diluted 1:5 with demi-water) and the samples were incubated for 5 min (RT), before the absorption was measured using a platereader (595 nm).

### Fluorescence microscopy to monitor osteogenic differentiation

3D cell cultures were fluorescently stained and imaged weekly during 5 to 7 weeks and at days 64 and 84 of differentiation. The gels were incubated for at least 2h with CNA-OG488 (0.167μM in OIM, 488 nm, TU/e (Aper et al., 2014)) in the incubator. Additionally, calcein blue (0.1 mg/mL, 405 nm (M1255)) and cell mask deep red (5 μg/mL, 630 nm (ThermoFisher, C10046)) were added to the staining medium for 2h. Next, the staining solution was removed, and the gels were washed twice with PBS (5 min, 37°C), before fresh medium was added. Using an LSM900 upright (Carl Zeiss Microscopy GmbH, Germany), overview tile scans were made of the complete gel (objective: 5x C Epiplan-APOCHROMAT/0.2 DIC M27, z-resolution=6.11μm, averaging=2, SR-4Y). Additionally, z-stacks were recorded of one region of interest per gel (objective 10x C Epiplan-APOCHROMAT/0.4 DIC M27, z-resolution=1.48 μm, averaging=2, SR-4Y). After imaging, the medium was replaced, and the cells were placed back in the incubator. For each experiment, the imaging settings were kept the same during the full duration of the experiment.

For live/dead staining Calcein-AM (500 nM) and propidium iodide (1 μg/mL) were diluted in OIM. The cells were incubated for 30 min (37°C) with the staining medium and analyzed immediately after staining using a LSM900 upright microscope to record z-stacks with 5x and 10x objectives as described above.

### Sample preparation for SEM

At day 49, two gels were washed with PBS after fluorescent imaging and fixed in 2% glutaraldehyde (in 0.1M Cacodylate buffer, pH = 7.4) for 90 min, while shaking (RT). After fixation, the gels were washed once quickly in cacodylate buffer and sequentially washed twice for 30 min (shaking, RT). Next, the gels were embedded in 4% ultra-low gelling temperature agarose (A5030, in MQ). The Ibidi slide was cooled on ice (30 min) to allow the agarose to gel. Afterward, the agarose blocks containing the cell cultures were removed from the well and stored in cacodylate buffer (4°C).

For post-fixation and staining, the samples were incubated in 2% OsO_4_/1.5% K_3_Fe(CN)_6_/2 mM CaCl_2_ in cacodylate buffer (0.1M) for 30 min on ice, followed by an additional incubation at room temperature (30 min, shaking). Next, the samples were washed in ddH_2_O (5x 5 min each) and incubated in thiocarbohydrazide (0.5% in MilliQ, 30 min, RT, shaking). The samples were washed in MilliQ (5x 5 min) and placed in 2% OsO_4_ (in MilliQ, 30 min, RT, shaking). After washing in MilliQ (5x 5 min), the samples were placed in 2% aqueous uranyl acetate (overnight, 4°C). Afterward, the samples were washed in ddH_2_O (5x 5 min) and stained in lead aspartate solution (pH 5.5) for 30 min (60°C). Next, the samples were washed in ddH_2_O (5x 5 min) and dehydrated on ice using an ascending ethanol series (50%, 70%, 90%, 3x 100% EtOH, 15 min per step, shaking) after which the samples were placed in acetone for 10 min (on ice). Sequentially, the samples were embedded in Durcupan by step-wise increasing the concentration of Durcupan in acetone to 100% Durcupan (25% 1.5h; 50% 1.5h; 75% overnight; 100%, twice 2.5h, 1.5h). The Durcupan was polymerized at 60°C for 48h.

### Sectioning for 3D Array Tomography SEM

One of the embedded cultures was sectioned for array tomography SEM (AT-SEM). Ribbons of serial ultrathin sections (400x200 μm, 80 nm thick, ∼250 sections) were made using a Leica Artos 3D Ultramicrotome (Leica AT4 knife). The ribbons were placed on ITO glass and coated with 5 nm carbon (Leica ACE600) and overview images (∼170x480 μm) were made of all sections using a Zeiss Sigma 300 SEM (50 nm/pixel, Atlas 5), after which higher resolution stacks were recorded (9 nm/pixel, 53x143 μm) from a selected region of interest.

Additionally, a section (200 nm thick) along the axial axis of the culture was prepared on ITO glass. The section was coated with 5nm carbon and an overview image was made using a Zeiss Sigma 300 SEM with Atlas 5 software (3kV, pixel size 15 nm), using the backscatter detector.

### SEM and 3D FIB/SEM

One embedded culture was sectioned along the longitudinal axis on a microtome until the core of the culture was reached. An overview of the sample was imaged with a Zeiss Sigma 300 SEM (3kV, pixel size 35 nm) and regions of interest were recorded (3kV, pixel size 10 nm). The sample was then coated with a gold sputter coater (Edwards) and transferred to a Zeiss Crossbeam 550 FIB/SEM.

At the regions of interest, Pt and C were sequentially deposited before a coarse trench was milled using a 30 kV/30 nA FIB probe and smoothened with a 30 kV/1.5 nA FIB probe. Parameters for serial sectioning imaging were set in Atlas 3D software (Atlas Engine v5.3.3) to allow imaging at a 10x10x10 nm voxel size, using a 30 kV/700 pA probe for serial FIB milling. InLens secondary and backscattered (grid -876V) electron microscopy images were simultaneously recorded (line average n=1, dwell time 0.7x4 μs) using an acceleration voltage of 2.0 kV and a probe current of 500 pA. After imaging, the ESB and InLens data were exported from Atlas as a combined file, using a 0.25 ratio of Inlens/ESB.

### Image analysis

Fluorescence images were analyzed in FIJI (version 1.52p). Using a macro (Supplementary Information B), maximum intensity projections (MIP) were made of the stitched tilescans (5x) and the brightness was adjusted for visualization purposes. For 10x images, a macro was used (Supplementary Information C) to prepare maximum intensity projections and perform orientation analysis with OrientationJ (σ=30 for cells; σ=12 for collagen, cubic spline) to extract color maps (OrientationJ Analysis (Püspöki et al., 2016)) as well as orientation histograms (OrientationJ Distribution (Rezakhaniha et al., 2012)). Orientation histograms were used in combination with the color maps to determine the quality of the analysis. In case no clear peak was present and the signal was too faint to determine reliable orientation color maps, the data were excluded from further analysis. The data from the orientation histograms (37 images from 4 experiments, containing data from 11 different gels) were then used to determine the peak maximum in Excel for the collagen as well as membrane orientation. The absolute difference between these values was used for analysis of the angles between the two components. To determine the full width at half maximum, the centers (35-145 degrees) of the histograms were transferred to Origin and fitted using a Lorentzian peak function.

### 3D Electron Microscopy Image Processing

AT-SEM data were aligned in Zeiss Atlas 5 and 210 sequential frames were exported. The frames were imported in Dragonfly ORS (Version 2022.2 for Windows, Comet Technologies Canada Inc. Montreal, Canada; software available at https://www.theobjects.com/dragonfly) and a U-NET++ network (U-NET++, Depth level: 5, Input Dimension: 3D) was trained for segmentation of cells in sequential steps, based on a total of 69 frames (∼1% of total voxels annotated). Voids were annotated based on a selected range from the image histogram (255-255). The inner areas in the voids ROI were filled (3D) before the ROI was smoothened (kernel 9, 3D).

The 3D FIB/SEM stacks were first processed in FIJI to sequentially remove shadows (subtract background, rolling ball radius 500 pixels), optimize the histogram (math, subtract, value = 75) and optimize the contrast (Local Contrast Enhancement (CLAHE), block size = 49, slope = 1.5).

After this, the stacks were visualized and segmented in Dragonfly ORS (Version 2022.2). For segmentation, one of the stacks was imported in the Dragonfly ORS segmentation wizard and 36 frames (0.11% of total voxels annotated) were manually annotated for background, collagen and cells (13% cells, 35.3% collagen, 51,7% background). A pre-trained Dragonfly neural network (YOLOv3, U-NET, Depth level: 5, Input Dimension: 2D, pre-trained on ORS500Kv01) was trained using these segmentations in 5 sequential steps, using increasing numbers of frames (from 5 to 36). To apply the model on the other full stacks, the initial trained model was optimized for the specific stack by importing the model in the Segmentation Wizard with the new stack and re-training on 5 (R16) or 8 (R13) manually assigned frames (R13: 0.15%/R16: 0.03% of total voxels annotated). For segmentation of cropped regions from all stacks, the initial model was used for segmentation using 2D or 3D (major vote) segmentation mode, while adjusting the brightness/contrast settings to correct for differences between the stacks. After segmentation, manual corrections and Boolean operations (e.g. fill holes) were applied to optimize the annotations. Next, the collagen was exported as ROI and used for orientation analysis in FIJI (OrientationJ, σ=3, cubic spline). For density measurement, ROIs of comparable sizes were selected that contained only collagen and the percentage of labeled voxels was copied from the Dragonfly characteristics.

## Supporting information

Supplementary Information B - Script MIPs - Language Image J Macro

Supplementary Information C - Script OriJ - Language Image J Macro

## Abbreviations

hMSCs: human mesenchymal stem cells
FIB/SEM: Focussed Ion Beam Scanning Electron Microscopy
AT-SEM: Array Tomography Scanning Electron Microscopy
ALP: Alkaline Phosphatase

## Acknowledgements

The authors thank Harrie Weinans and Ralph Sakkers from University Medical Center Utrecht for providing the hMSCs. The project was supported by an European Research Council (ERC) Advanced Investigator grant (H2020-ERC-2017-ADV-788982-COLMIN) to N.S. A.A. was also supported by a VENI grant from the Netherlands Scientific Organization NWO (VI.Veni.192.094).

## References

Akiva, A., Melke, J., Ansari, S., Liv, N., Meijden, R., Erp, M., Zhao, F., Stout, M., Nijhuis, W.H., Heus, C., Muñiz Ortera, C., Fermie, J., Klumperman, J., Ito, K., Sommerdijk, N., Hofmann, S., 2021. An Organoid for Woven Bone. Adv Funct Mater 31, 2010524.

Aper, S.J., van Spreeuwel, A.C., van Turnhout, M.C., van der Linden, A.J., Pieters, P.A., van der Zon, N.L., de la Rambelje, S.L., Bouten, C.V., Merkx, M., 2014. Colorful protein-based fluorescent probes for collagen imaging. PLoS One 9, e114983.

Blair, H.C., Larrouture, Q.C., Li, Y., Lin, H., Beer-Stoltz, D., Liu, L., Tuan, R.S., Robinson, L.J., Schlesinger, P.H., Nelson, D.J., 2017. Osteoblast Differentiation and Bone Matrix Formation In Vivo and In Vitro. Tissue Eng Part B Rev 23, 268–280.

Boraschi-Diaz, I., Wang, J., Mort, J.S., Komarova, S.V., 2017. Collagen Type I as a Ligand for Receptor-Mediated Signaling. Frontiers in Physics 5.

Budyn, E., Gaci, N., Sanders, S., Bensidhoum, M., Schmidt, E., Cinquin, B., Tauc, P., Petite, H., 2018. Human Stem Cell Derived Osteocytes in Bone-on-Chip. MRS Advances 3, 1443–1455.

Buss, D.J., Kröger, R., McKee, M.D., Reznikov, N., 2022. Hierarchical organization of bone in three dimensions: A twist of twists. Journal of Structural Biology: X 6, 100057.

de Jonge, N., Kanters, F.M., Baaijens, F.P., Bouten, C.V., 2013. Strain-induced collagen organization at the microlevel in fibrin-based engineered tissue constructs. Ann Biomed Eng 41, 763–774.

de Wildt, B.W.M., Ansari, S., Sommerdijk, N.A.J.M., Ito, K., Akiva, A., Hofmann, S., 2019. From bone regeneration to three-dimensional in vitro models: tissue engineering of organized bone extracellular matrix. Current Opinion in Biomedical Engineering 10, 107–115.

Fisher, K.E., Sacharidou, A., Stratman, A.N., Mayo, A.M., Fisher, S.B., Mahan, R.D., Davis, M.J., Davis, G.E., 2009. MT1-MMP- and Cdc42-dependent signaling co-regulate cell invasion and tunnel formation in 3D collagen matrices. J Cell Sci 122, 4558–4569.

Fratzl, P., Weinkamer, R., 2007. Nature’s hierarchical materials. Progress in Materials Science 52, 1263–1334.

Golub, E.E., Boesze-Battaglia, K., 2007. The role of alkaline phosphatase in mineralization. Current Opinion in Orthopaedics 18, 444–448.

Guidry, C., Grinnell, F., 1987. Contraction of Hydrated Collagen Gels by Fibroblasts: Evidence for Two Mechanisms by which Collagen Fibrils are Stabilized. Collagen and Related Research 6, 515–529.

Husch, J.F.A., Coquelin, L., Chevallier, N., Tiemessen, D., Oosterwijk, E., van Rheden, R., Woud, C., Vossen, J., Leeuwenburgh, S.C.G., van den Beucken, J.J.J.P., 2023. Comparison of Osteogenic Capacity and Osteoinduction of Adipose Tissue-Derived Cell Populations. Tissue Eng Part C Methods 29, 216–227.

Iordachescu, A., Amin, H.D., Rankin, S.M., Williams, R.L., Yapp, C., Bannerman, A., Pacureanu, A., Addison, O., Hulley, P.A., Grover, L.M., 2018. An In Vitro Model for the Development of Mature Bone Containing an Osteocyte Network. Adv Biosyst 2, 1700156.

Jaiswal, N., Haynesworth, S.E., Caplan, A.I., Bruder, S.P., 1998. Osteogenic differentiation of purified, culture-expanded human mesenchymal stem cells in vitro. Journal of Cellular Biochemistry 64, 295–312.

Kassem, M., Rungby, J., Mosekilde, L., Eriksen, E.F., 1992. Ultrastructure of human osteoblasts and associated matrix in culture. APMIS 100, 490–497.

Katsimbri, P., 2017. The biology of normal bone remodelling. Eur J Cancer Care (Engl) 26.

Kim, D.-H., Ewald, A.J., Park, J., Kshitiz Kwak, M., Gray, R.S., Su, C.-Y., Seo, J., An, S.S., Levchenko, A., 2018. Biomechanical interplay between anisotropic re-organization of cells and the surrounding matrix underlies transition to invasive cancer spread. Scientific Reports 8, 14210.

Kim, J., Adachi, T., 2019. Cell Condensation Triggers the Differentiation of Osteoblast Precursor Cells to Osteocyte-Like Cells. Front. Bioeng. Biotech. 7.

Kim, J., Ishikawa, K., Sunaga, J., Adachi, T., 2021. Uniaxially fixed mechanical boundary condition elicits cellular alignment in collagen matrix with induction of osteogenesis. Scientific Reports 11, 9009.

Kundu, A.K., Khatiwala, C.B., Putnam, A.J., 2009. Extracellular matrix remodeling, integrin expression, and downstream signaling pathways influence the osteogenic differentiation of mesenchymal stem cells on poly(lactide-co-glycolide) substrates. Tissue Eng Part A 15, 273–283.

Lange, J.R., Fabry, B., 2013. Cell and tissue mechanics in cell migration. Experimental Cell Research 319, 2418–2423.

Lee, H.M., Seo, S.-R., Kim, J., Kim, M.K., Seo, H., Kim, K.S., Jang, Y.-J., Ryu, C.J., 2020. Expression dynamics of integrin α2, α3, and αV upon osteogenic differentiation of human mesenchymal stem cells. Stem Cell Research & Therapy 11, 210.

Lu, P., Lu, Y., 2021. Born to Run? Diverse Modes of Epithelial Migration. Front. Cell. Dev. Biol. 9, 704939.

Lu, Y., Kamel-El Sayed, S.A., Wang, K., Tiede-Lewis, L.M., Grillo, M.A., Veno, P.A., Dusevich, V., Phillips, C.L., Bonewald, L.F., Dallas, S.L., 2018. Live Imaging of Type I Collagen Assembly Dynamics in Osteoblasts Stably Expressing GFP and mCherry-Tagged Collagen Constructs. J Bone Miner Res 33, 1166–1182.

Magal, R.A., Reznikov, N., Shahar, R., Weiner, S., 2014. Three-dimensional structure of minipig fibrolamellar bone: Adaptation to axial loading. Journal of Structural Biology 186, 253–264.

Matsumoto, T., Sasaki, J., Alsberg, E., Egusa, H., Yatani, H., Sohmura, T., 2007. Three-dimensional cell and tissue patterning in a strained fibrin gel system. PLoS One 2, e1211.

Meesuk, L., Suwanprateeb, J., Thammarakcharoen, F., Tantrawatpan, C., Kheolamai, P., Palang, I., Tantikanlayaporn, D., Manochantr, S., 2022. Osteogenic differentiation and proliferation potentials of human bone marrow and umbilical cord-derived mesenchymal stem cells on the 3D-printed hydroxyapatite scaffolds. Sci Rep 12, 19509.

Miller, A.E., Hu, P., Barker, T.H., 2020. Feeling Things Out: Bidirectional Signaling of the Cell–ECM Interface, Implications in the Mechanobiology of Cell Spreading, Migration, Proliferation, and Differentiation. Adv Healthc Mater 9, 1901445.

Miron, R.J., Zhang, Y.F., 2012. Osteoinduction: A Review of Old Concepts with New Standards. Journal of Dental Research 91, 736–744.

Nasello, G., Alaman-Diez, P., Schiavi, J., Perez, M.A., McNamara, L., Garcia-Aznar, J.M., 2020. Primary Human Osteoblasts Cultured in a 3D Microenvironment Create a Unique Representative Model of Their Differentiation Into Osteocytes. Front Bioeng Biotechnol 8, 336.

Pamonag, M., Hinson, A., Burton, E.J., Jafari, N., Sales, D., Babcock, S., Basha, R., Hu, X., Kubow, K.E., 2022. Individual cells generate their own self-reinforcing contact guidance cues through local matrix fiber remodeling. PLoS One 17, e0265403.

Pennings, I., van Dijk, L.A., van Huuksloot, J., Fledderus, J.O., Schepers, K., Braat, A.K., Hsiao, E.C., Barruet, E., Morales, B.M., Verhaar, M.C., Rosenberg, A.J.W.P., Gawlitta, D., 2019. Effect of donor variation on osteogenesis and vasculogenesis in hydrogel cocultures. Tissue Engineering and Regenerative Medicine 13, 433–445.

Püspöki, Z., Storath, M., Sage, D., Unser, M., 2016. Transforms and Operators for Directional Bioimage Analysis: A Survey. Adv. Anat. Embryol. Cell Biol. 219, 69–93.

Rezakhaniha, R., Agianniotis, A., Schrauwen, J.T.C., Griffa, A., Sage, D., Bouten, C.V.C., van de Vosse, F.N., Unser, M., Stergiopulos, N., 2012. Experimental investigation of collagen waviness and orientation in the arterial adventitia using confocal laser scanning microscopy. Biomechanics and Modeling in Mechanobiology 11, 461–473.

Reznikov, N., Shahar, R., Weiner, S., 2014. Bone hierarchical structure in three dimensions. Acta Biomater 10, 3815–3826.

Rodriguez-Pascual, F., Slatter, D.A., 2016. Collagen cross-linking: insights on the evolution of metazoan extracellular matrix. Scientific Reports 6, 37374.

Sasaki, J.-I., Matsumoto, T., Egusa, H., Nakano, T., Ishimoto, T., Sohmura, T., Yatani, H., 2010. In vitro engineering of transitional tissue by patterning and functional control of cells in fibrin gel. Soft Matter 6.

Sasaki, J., Matsumoto, T., Imazato, S., 2015. Oriented bone formation using biomimetic fibrin hydrogels with threedimensional patterned bone matrices. Journal of Biomedical Materials Research Part A 103, 622–627.

Stains, J.P., Civitelli, R., 2005. Cell-cell interactions in regulating osteogenesis and osteoblast function. Birth Defects Research Part C: Embryo Today: Reviews 75, 72–80.

Tenenbaum, H.C., 1981. Role of Organic Phosphate in Mineralization of Bone in vitro. Journal of Dental Research 60, 1586–1589.

Tenenbaum, H.C., Heersche, J.N.M., 1982. Differentiation of osteoblasts and formation of mineralized bone in vitro. Calcified Tissue International 34, 76–79.

Thrivikraman, G., Athirasala, A., Gordon, R., Zhang, L., Bergan, R., Keene, D.R., Jones, J.M., Xie, H., Chen, Z., Tao, J., Wingender, B., Gower, L., Ferracane, J.L., Bertassoni, L.E., 2019. Rapid fabrication of vascularized and innervated cell-laden bone models with biomimetic intrafibrillar collagen mineralization. Nat Commun 10, 3520.

Vogel, W., Gish, G.D., Alves, F., Pawson, T., 1997. The Discoidin Domain Receptor Tyrosine Kinases Are Activated by Collagen. Molecular Cell 1, 13–23.

Weiner, S., Wagner, H.D., 1998. THE MATERIAL BONE: Structure-Mechanical Function Relations. Annu Rev Mater Sci 28, 271–298.

Westhauser, F., Karadjian, M., Essers, C., Senger, A.-S., Hagmann, S., Schmidmaier, G., Moghaddam, A., 2019. Osteogenic differentiation of mesenchymal stem cells is enhanced in a 45S5-supplemented β-TCP composite scaffold: an in-vitro comparison of Vitoss and Vitoss BA. PLoS One 14, e0212799.

Yang, H., Sun, L., Cai, W., Gu, J., Xu, D., Deb, A., Duan, J., 2020. DDR2, a discoidin domain receptor, is a marker of periosteal osteoblast and osteoblast progenitors. Journal of Bone and Mineral Metabolism 38, 670–677.

Zernik, J., Twarog, K., Upholt, W.B., 1990. Regulation of alkaline phosphatase and alpha2(I) procollagen synthesis during early intramembranous bone formation in the rat mandible. Differentiation 44, 207–215.

