## Supplementary A - Figures & Table for "Osteoblast-Induced Collagen Alignment in a 3D *in vitro* Bone Model"

**Supplementary Figures**

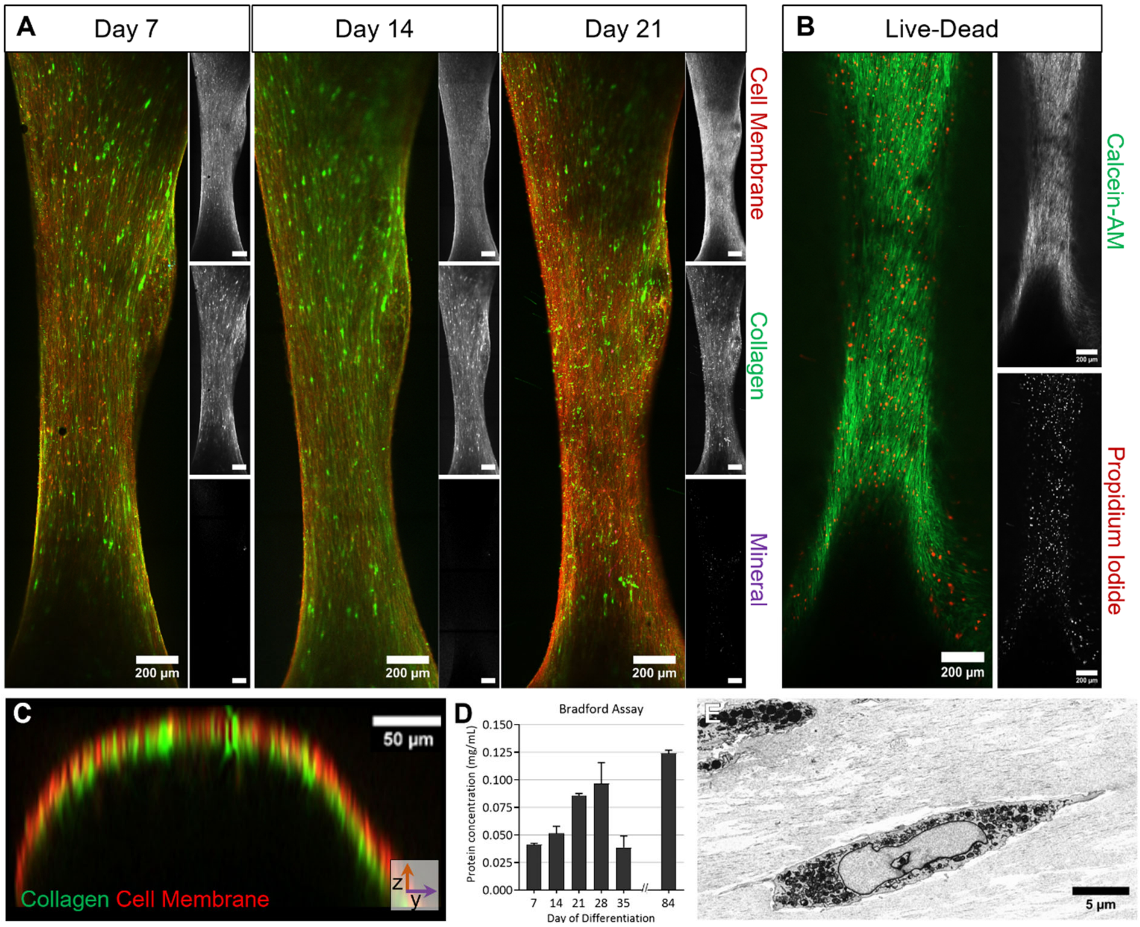


***Supplementary Figure A.1***

*A) The cellular distribution (red), collagen production (green), and mineral deposition (magenta) as visualized by overview fluorescence images at 7/14/21 days of differentiation. Single channel images are shown in black and white next to the merged image. Scalebars: 200 µm. B) Live-Dead staining showing living cells by calcein-AM (green) and dead cells by propidium iodide (red) staining after 21 days of differentiation. Single channel images are shown in black and white. Scalebar: 200 µm. C) Reconstruction of axial plane (XZ) from fluorescence microscopy z-stack shows that only the outer regions (periphery and partially bulk) can be visualized. Collagen (green) and cell membranes (red). Scalebar: 50 µm. D) Bradford assay at different times of differentiation, used for normalization of the ALP assay. E) High-resolution (9 nm/pixel) SEM image showing cell morphology and organelles of cells surrounded by collagen. Scalebar: 5 µm.*

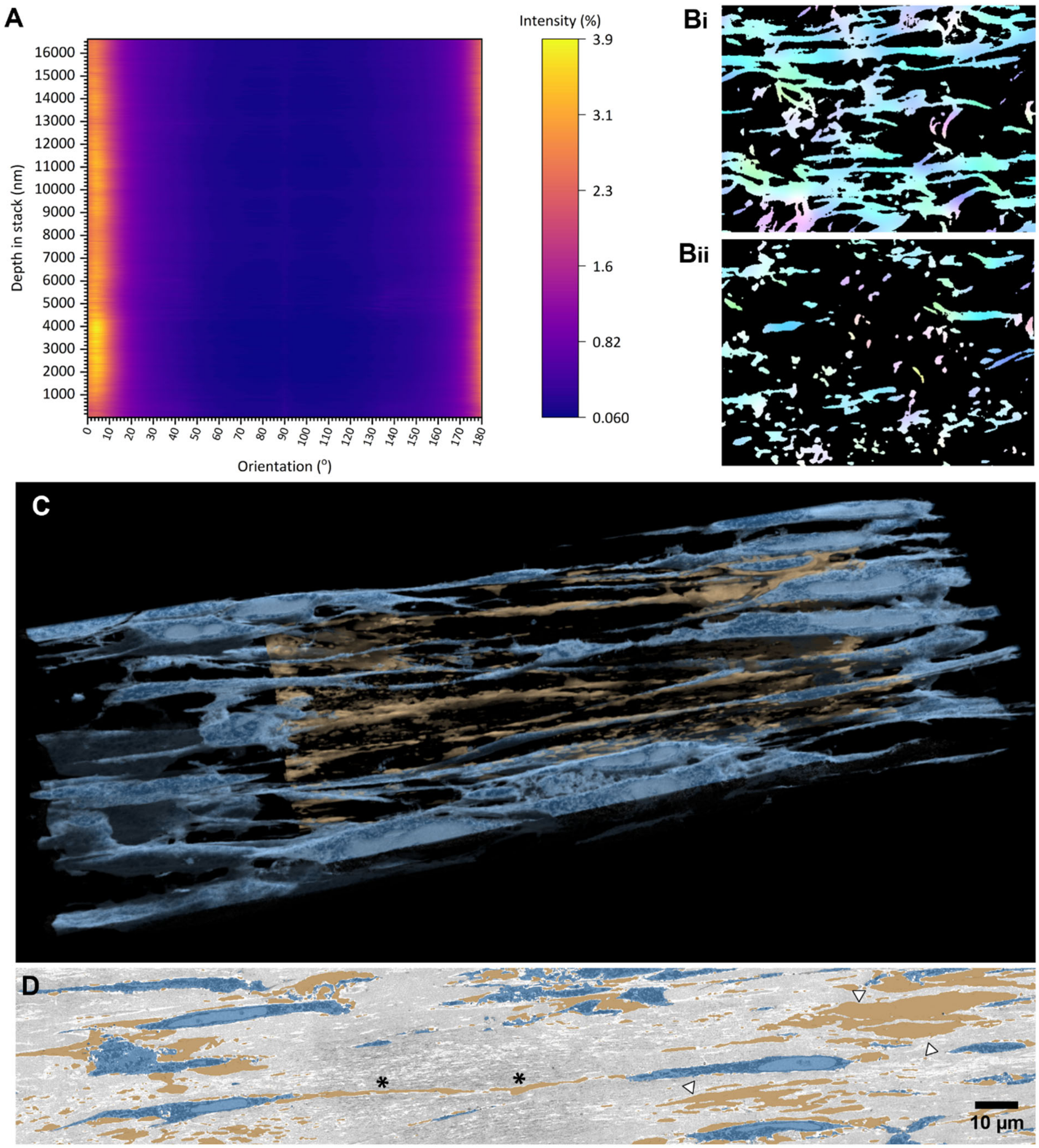


***Supplementary Figure A.2***

*A) Heatmap representation of the orientation histograms throughout a stack in the bulk region. The intensity (%) indicates the percentage of pixels with a certain orientation. B) Color-based orientation analysis of the segmented collagen in regions with varying collagen densities. In the bulk region closer to the periphery but without cells collagen is clearly still aligned. Collagen in the bulk located closer to the core region shows lower density and lower degrees of orientation. C) 3D reconstruction of cells segmented from AT-SEM imaging in the core region and surrounded by tunnels and voids (brown) without cellular or protein structures. D) SEM image in the core region showing smaller tunnels in regions with fibrin mixed with collagen (*) compared to regions without collagen (▼). Scalebar: 10 μm.*

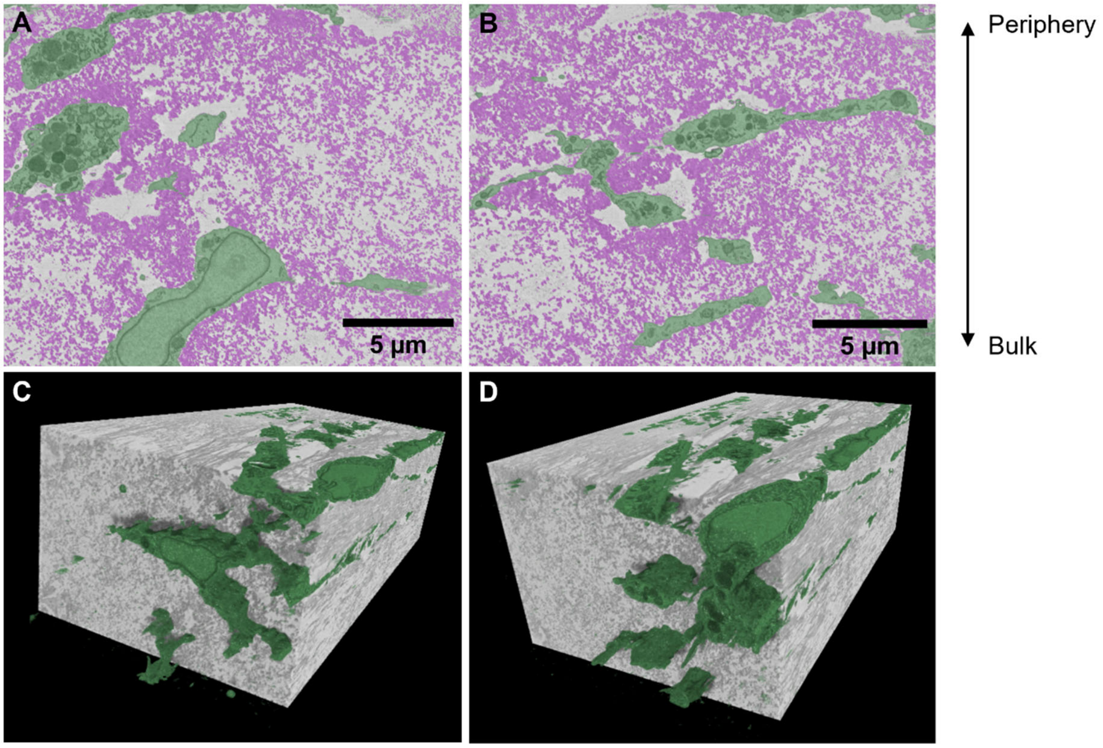


***Supplementary Figure A.3***
*A/B) Additional axial planes of cells (green) in the dense collagen region (magenta) close to the periphery, showing long protrusions next to thicker cell-bodies. C/D) 3D Reconstructions of cells (green) in the dense collagen region.*

**Supplementary Table A.1 – Collagen Densities in Variable Regions of the Bulk**

| Visual Characterization of Collagen Density | Labeled Voxels in Volume (%) *1 | Analyzed Total Volume ($nm^3$) |
|---|---|---|
| **Dense** | 67 | 82.436.942.000 *2 |
| **Intermediate** | 34 | 581.605.110.000 |
| **Low** | 15 | 656.271.000.000 |

*1 Collagen density was determined based on voxel labeling of automated segmentation in selected volumes in which no cells were present.
*2 Sum of 3 regions; percentage of labeled voxels was calculated as the average of the labeling in these three regions.
